# Fast principal components analysis reveals convergent evolution of ADH1B gene in Europe and East Asia

**DOI:** 10.1101/018143

**Authors:** Kevin J. Galinsky, Gaurav Bhatia, Po-Ru Loh, Stoyan Georgiev, Sayan Mukherjee, Nick J. Patterson, Alkes L. Price

## Abstract

Searching for genetic variants with unusual differentiation between subpopulations is an established approach for identifying signals of natural selection. However, existing methods generally require discrete subpopulations. We introduce a method that infers selection using principal components (PCs) by identifying variants whose differentiation along top PCs is significantly greater than the null distribution of genetic drift. To enable the application of this method to large data sets, we developed the FastPCA software, which employs recent advances in random matrix theory to accurately approximate top PCs while reducing time and memory cost from quadratic to linear in the number of individuals, a computational improvement of many orders of magnitude. We apply FastPCA to a cohort of 54,734 European Americans, identifying 5 distinct subpopulations spanning the top 4 PCs. Using the PC-based test for natural selection, we replicate previously known selected loci and identify three new genome-wide significant signals of selection, including selection in Europeans at the ADH1B gene. The coding variant rs1229984*T has previously been associated to a decreased risk of alcoholism and shown to be under selection in East Asians; we show that it is a rare example of independent evolution on two continents. We also detect new selection signals at IGFBP3 and IGH, which have also previously been associated to human disease.

## Introduction

Searching for genetic variants with unusual differentiation between populations is an established approach for identifying signals of natural selection^1–6^. We and others have employed this approach to identify signals of selection in a wide range of settings, informing our understanding of genes under evolutionary adaptation^7–24^. Examples includes genes linked to lactase persistence^9,11^, starch hydrolysis^12^, fatty acid decomposition^24^, red blood cell abundance^17^, hypoxia response^18^, alcoholism^14^, kidney disease^21^, malaria^7,13,19,23^, HIV/AIDS^16^, autoimmune disease^20^, cancer^19^, cystic fibrosis^8^ and hypertension^23^. However, the signals of selection identified thus far may represent “only the tip of the iceberg^25^”, implying that further research on selection will provide additional insights about human disease. Unlike extended haplotype homozygosity (EHH) or allele frequency spectrum based tests for selection, the population differentiation approach is able to detect older selection events and selection on standing variation^1,3^. In addition, signals of selection detected using population differentiation can flag stratified genetic variants that are susceptible to false-positive associations in genome-wide association studies^15^.

Previous work on detecting selection using population differentiation has focused on methods that evaluate deviations from genome-wide patterns of genetic drift between discrete populations, such as Locus-Specific Branch Length (LSBL)^6^, Population Branch Statistic (PBS)^17^ and TreeSelect^19^. The population differentiation approach has greatest power when comparing very closely related populations with very large sample size^19^. The increasing availability of very large population cohorts for genetic analysis provides strong prospects for analyzing subtle differences in ancestry in large sample sizes, but raises the challenge of how to select subpopulations to compare; a population cohort with a single continental ancestry may be better represented by continuous clines rather than discrete clusters^26–28^, and/or may contain a large number of discrete subpopulations corresponding to a large number of possible population comparisons^29,30^. Principal components analysis (PCA)^26,31^ offers an appealing alternative to model-based clustering methods^32,33^ for modeling human genetic diversity, and has been applied to infer population structure in many settings^27,28,31,34–40^. One advantage of PCA is that results for top PCs are not sensitive to the number of PCs analyzed, whereas results of model-based clustering methods often vary with the number of clusters. Another advantage of PCA is its low computational cost, as top PCs can be inferred in time only linear in the number of samples by drawing upon recent advances in random matrix theory^41–43^, implemented in the FastPCA software that we introduce here. We thus developed a test for selection that uses the SNP weights from PCA to calculate the differentiation of each locus along top PCs. Specifically, the squared correlation of each SNP to a PC, rescaled to account for genetic drift, follows a chi-square (1 d.o.f.) distribution under the null hypothesis of no selection. Our PC-based test produces a *p*-value at each locus and is able to detect novel signals at genome-wide significance, a key consideration in genome scans for selection^19^.

We ran FastPCA on 54,734 individuals of European descent from the Genetic Epidemiology Research on Adult Health and Aging (GERA) cohort; FastPCA required only 57 minutes of compute time and 2.6GB of RAM for this analysis, orders of magnitude better than any other publicly available software. We detected evidence of population structure along the top 4 PCs, which separated samples into several subpopulations. Using our PC-based test for selection, we replicate previously known selected loci *LCT* [MIM 603202], *HLA* [MIM 142800], *OCA2* [MIM 611409] and *IRF4* [MIM 601900] and identify three new signals of selection at *IGH* [MIM 147100], *IGFBP3* [MIM 146732] and *ADH1B* [MIM 103720]. The signal in *ADH1B* at coding variant rs1229984 has previously been associated to alcoholism^44–47^ and shown to be under selection in East Asians^14,46,48,49^; we show that it is a rare example of independent evolution on two continents^11,12^.

## Methods

### Overview of methods

We first describe the FastPCA algorithm, which is an implementation of the *blanczos* method from Rokhlin *et al*.^41–43^. As with our previous work on PCA^26,31^, FastPCA makes use of existing computational literature and does not contain any new computational ideas; nonetheless, we anticipate that the software will be widely used, since to our knowledge it is the only publicly available software for computing top PCs on genetic data in linear time. The algorithm generalizes the method of power iteration^50^, a technique to estimate the largest eigenvalue and corresponding eigenvector of a matrix. Multiplying a random vector by a square matrix projects that vector onto the eigenvectors of that matrix and then scales it according the respective eigenvalues of that matrix. After repeating, the projection along the eigenvector with the largest eigenvalue grows fasters than the rest and the repeated matrix by vector product converges to this eigenvector. Additional eigenvectors can be found by repeating this process and orthogonalizing to previously-found PCs. The *blanczos* method improves on this method by initially estimating more PCs than ultimately desired. The original estimates are perturbed from the true PCs, but this missing variation is captured by estimating the extra PCs. The genotype matrix is then projected onto this set of eigenvectors, reducing its dimension while preserving the variation along the top PCs. Traditional PCA methods are applied to this reduced matrix to find accurate estimates of the top PCs of the original matrix.

We next describe our PC-based selection statistic, which generalizes a previous selection statistic developed for discrete populations^19^. We detect unusual allele frequency differences along inferred PCs by making use of the fact that the squared correlation of each SNP to a PC, rescaled to account for genetic drift (Selection statistic), follows a chi-square (1 d.o.f.) distribution under the null hypothesis of no selection. We have released open-source software implementing the FastPCA algorithm and PC-based selection statistic (see Web Resources).

### FastPCA algorithm

We are given an input *M × N* genotype matrix ***X***, where *M* is the number of SNPs and *N* is the number of individuals (e.g. each row is a SNP, each column is a sample). Each entry in this matrix takes its values from {0,1,2} indicating the count of variant alleles for a sample at a SNP. From this matrix we can generate the normalized genomic matrix 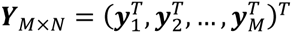 where each row ***y_i_*** has approximately mean 0 and variance 1 for SNPs in Hardy-Weinberg equilibrium.

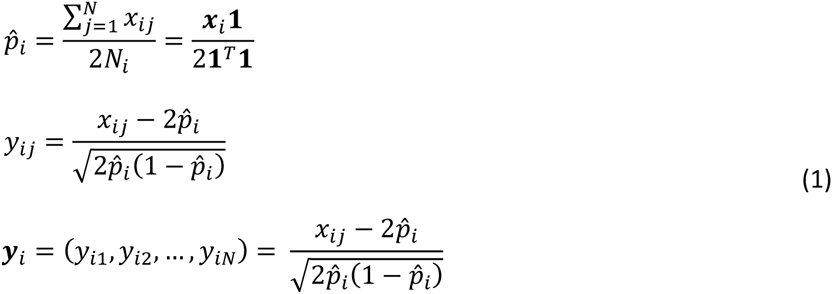

Here, ***x****_i_* is the row vector of genotypes for SNP *i* and ***y****_i_* is the normalized row vector. *x_ij_* and *y_ij_* are the genotype/normalized genotype at SNP *i* for sample *j. N_i_* is the number of valid genotypes at SNP *i*. All this is used to calculate 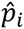, the sample allele frequency for SNP *i*, which is used to normalize the genotypes. In practice, the genotype matrix is normalized through the use of a lookup table mapping from genotypes (stored as 0, 1 or 2 copies of the alternate allele, or missing data) to normalized genotypes (using the above formula, with missing data having a normalized value of 0).

We are seeking the top *K* PCs for the normalized genomic matrix ***Y***. Traditional PCA algorithms compute the PCs by performing the eigendecomposition of the genetic relationship matrix (*GRM* = ***Y****^T^****Y****/M*), a costly procedure which returns all the principal components. FastPCA, which makes use of recent advances in random matrix theory^41–43^, speeds this process up by only approximating the top *K* PCs.

FastPCA is seeded with a random *N × L* matrix ***G***_0_ composed of values drawn from a standard Gaussian distribution. *L* affects the accuracy of the result and *L* should be greater than *K.* For *K* = 10, *L* = 20 is a good choice. Then, for *I* iterations, we calculate ***H****_i_* = ***Y*** *×* ***G****_i_* and ***G****_i_*+_1_ = ***Y****^T^ ×* ***H****_i_/M,* where the ***H****_i_*s are *M × L* matrices and ***G****_i_*s are *N × L* matrices like ***G***_0_. In simulated samples with discrete populations, *I = 3* was sufficient, but in real datasets, *I* = 10 was found to provide accurate results.

After the iterative step completes, we stack the ***H****_i_* matrices to produce the matrix ***H****_M×_*_(_*_I+_*_1)_*_L_* = (***H***_0_, ***H***_1_, …,***H****_I_*) and the singular value decomposition of matrix ***H*** is taken: 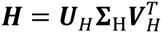. ***U****_H_* is a low-rank approximation to the column-space of ***Y*** with dimension *M ×* (*I + 1*)*L,* where 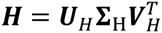. ***Y*** is then projected onto ***U****_H_* to produce ***T***_(_*_I+_*_1)_*_L×N_* = ***U****_H_****Y***. The SVD of 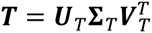 can be computed efficiently and approximates the SVD of ***Y*** since 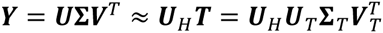. For the PCA, we are only interested in the left *K* columns of ***V****_T_* and the first *K* entries along the diagonal of ***Σ****_T_*.

FastPCA runs in linear time and memory relative to *M* and *N.* There are *0*(*I*) matrix multiplications where each multiplication takes *O*(*MNL*) time. Then, the SVD of *H* takes *0*(*MI^2^L^2^*) and the SVD of *T* takes *0*(*NI^2^L^2^*) time. Taking *I* and *L* to be constants, the overall running time simplifies to *O*(*MN*).

### Selection statistic

We first consider the simple case of an ancestral population that split into two extant populations with genetic distance *F_ST_*. We consider the allele frequencies at SNP *i* for the ancestral population (*p_i_*) and the two extant populations (*p_i_*_1_ and *p_i_*_2_). If there is no selection and SNPs are randomly ascertained, *p_i_*_1_−*p_i_*_2_ has expectation 0 (because allele frequencies can drift either up or down in each population) and variance 2*pi*(1 *− p_i_*)*F_ST_*^51^. In the case where *p_i_* is not close to 0 or 1 and *F_ST_* is small, the distribution of this difference approximately follows a normal distribution:

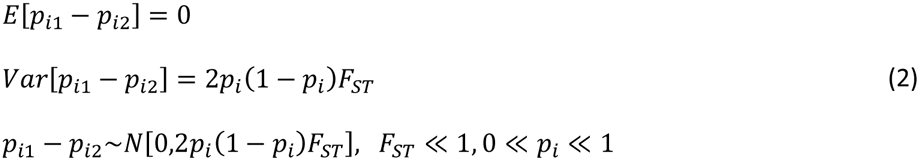

In practice, we do not have access to either the ancestral allele frequency or the extant population allele frequencies. Instead, we have sample allele frequencies for the two extant populations, 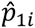 and 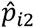. Assuming a large enough sample size from each population (*N*_1_ and *N_2_*) and that the true population allele frequency is not close to 0 or 1, these sample allele frequency estimates approximately follow a normal distribution with respect to the true allele frequencies. If we additionally assume that the ancestral allele frequency can be approximated by averaging the sample allele frequencies and that the true population allele frequencies are not that different, the sample allele frequency difference also follows a normal distribution^13,15,19^:

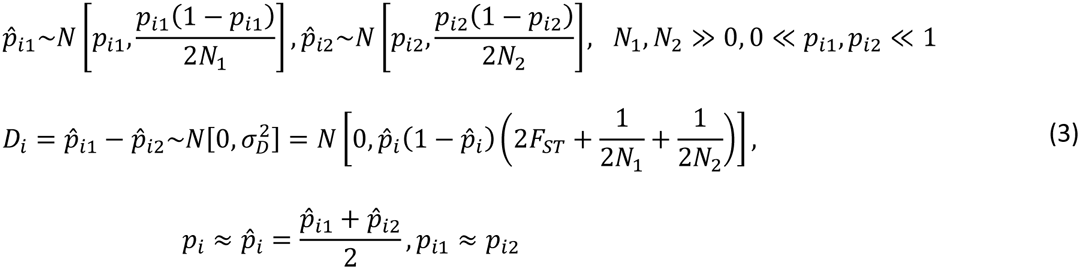

Below, we build the intuition behind our PC-based statistic by rewriting the discrete-population statistic using vector notation, then extending this statistic to individuals with fractional ancestries, and then to continuous-valued PCs.

In the case with two discrete populations, we define a vector ***α*** where *α_j_* indicates the ancestry in population 1 (e.g. *α_j_ = 1* if sample *j* is in population 1 and 0 if sample *j* is in population 2). *D_i_* can be rewritten as:

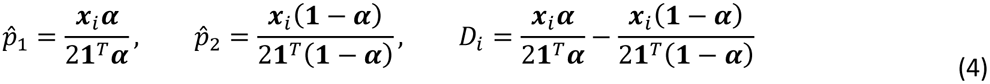

If we run PCA on this sample, we would ideally get an eigenvector ***v*** that has value *v*_1_ for individuals in population 1 and *−v*_2_ for individuals in population 2, where (since ***v****^T^***1** = *0,* ***v****^T^****v*** = 1)

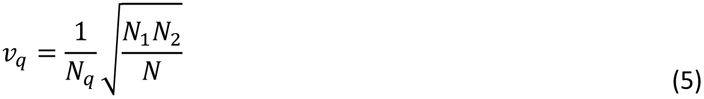

In this case, *D_i_* can be rewritten as:

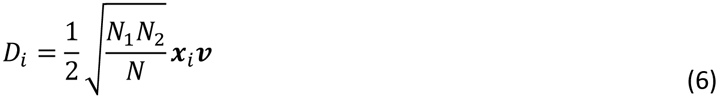

In the limiting case where *F_ST_* approaches 0, the statistic becomes:

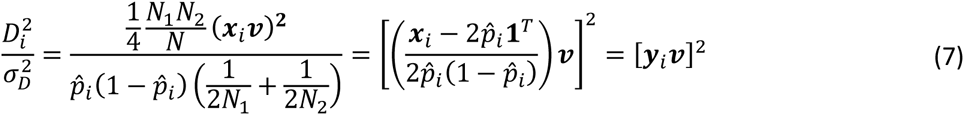

Thus, the square of the SNP weight follows a chi-square 1-d.o.f. distribution in the case where *F_ST_ →* 0. In the case where *F_ST_ ≠* 0, then the scaling parameter has to be changed, but *D_i_* still follows a normal distribution.

In the case with fractional ancestry (*α_j_ ∈* [0,1]), 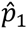, 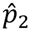 and *D_i_* can still be estimated using equation (4). The individual 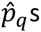 will still asymptotically follow a normal distribution (because of the Lyapunov central limit theorem^52^), but will be correlated due to individuals with fractional ancestry contributing to both estimates. Thus, *D_i_* will still follow a normal distribution, but the variance of equation (3) will not hold.

Now consider the case where we do not have fractional ancestries, but rather an eigenvector that separates individuals along some axis of variation. (We assume that extreme outlier individuals detected by PCA have been removed^26^, as PCs dominated by such outliers may violate normality assumptions.) We can treat the eigenvector as a linear transformation of the ancestry vector:

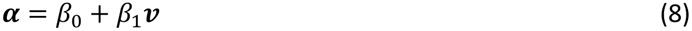

Substituting these values into (4), we find:

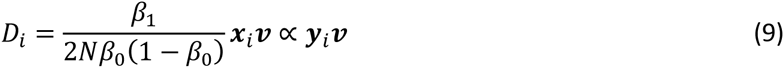

Thus, our new selection statistic *D_i_* is based on the dot product of the normalized genotypes and the eigenvector. Since the variance of *D_i_* is not known, it will need to be rescaled in order to follow a *N*(0,1^2^) distribution.

If we are operating on the same set of SNPs that we used for PCA, then the rescaling of ***y****_i_****v*** is straightforward. Because PCA is the same as SVD, we see that:

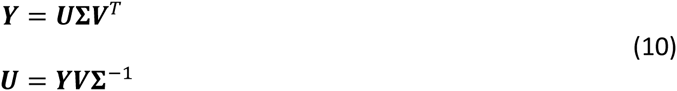

Here, ***V*** contains the right singular vectors which are equivalent to the PCs, ***U*** contains the left singular vectors which are rescaled SNP weights and ***Σ*** contains the singular values which are the square roots of the eigenvalues of the GRM. ***V*** and ***U*** are unitary, so the columns of ***U*** are guaranteed to have a norm of 1. Multiplying ***U*** by 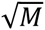 will then produce a properly normalized vector of differences (*D*_1_, *D*_2_ …, *D*_M_)*^T^* In other words:

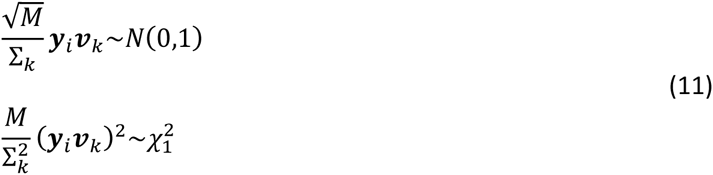

In the case of non-random SNP ascertainment and non-random choice of reference and variant allele, the expectation of *D_i_* may be non-zero. However, if we randomly flip the reference and variant alleles in such a situation, the resulting principal components and values of *D_i_* remain unchanged up to a factor of −1 and the expectation of *D_i_* becomes 0. As a result, even if there are systematically positive or negative SNP loadings, 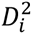 still follows a chi-square 1-d.o.f distribution.

In the case where we are computing selection statistics on a different set of SNPs than the one for which we computed PCs, then the above property is not guaranteed to hold. Specifically, inflation can occur if SNPs with higher differentiation tend to have higher LD, which can occur as a consequence of true selection signals^53^.

One assumption underlying the statistic is that the true minor allele frequency is not extremely small, otherwise the assumption of normality will not hold^19^. For this reason, the selection statistic was only computed for those SNPs containing minor allele frequency greater than 1% in our sample.

### Simulation framework

Genotypes were simulated at *M* independent SNPs and *N* independent individuals in four steps:

1. The ancestral allele frequency (*p_i_*) for a given SNP *i* was sampled from a *Uniform*(0.05,0.95) distribution.
2. Allele frequencies for *Q* populations (***P****_i_* = (*p_ii_,p_i2_, …,p_iq_*)*^T^*) were generated by simulating random drift (see below).
3. Admixture (***α****_j_*) for individual *j* was sampled from a *Dirichlet*(***a***) distribution.
4. Genotype *g_ij_* was sampled from a 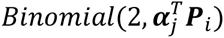 distribution.

Population allele frequencies were generated by simulating random drift in *Q* populations of fixed size *N_e_* for *τ* generations and stored in *Q ×* 1 vector ***P****_i_* = (*p_i_*_1_*, p_i2_,*..., *p_iq_*)*^T^*. The number of alternate alleles *z_iqt_* at SNP *i* in population *q* at generation *t* were sampled from a *Binomial*(*2N_e_,p_i,q,t−_*_1_) distribution, where *p_iq_*_0_ is the ancestral allele frequency *p_i_*. The population allele frequency at this generation was then calculated as 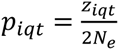. For most simulations, population allele frequency simulations were run for *τ* = 200 total generations and population size *N_e_* was calculated for a target *F_sT_* by using the formula 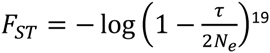. For *F_ST_* ≈ 0.1, 0.01 and 0.001, *N_e_ = 1k, 10k* and *100k* respectively. To detect the effect of population bottlenecks at the same level of *F_ST_*, simulations were also run for *τ* = 20 and *N_e_* = 100, *1k* and *10k,* again producing populations with genetic distance *F_sT_* ≈ 0.1, 0.01 and 0.001. Most simulations were run with two populations, but we also simulated 5 populations with a phylogenetic structure as follows. We set *N_e_ = 10k* and *τ* = 200 for populations 1 and 2, and *τ* = 180 for an intermediary ancestral population of populations 3, 4 and 5, yielding allele frequency 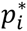. This was then fed back into the random drift model for an additional 20 generations for populations 3, 4 and 5. The pairwise genetic distance between populations 3, 4 and 5 is *F_ST_* ≈ 0.001 while the genetic distance between any other pair of populations is *F_ST_* ≈ 0.01.

We also considered simulations with admixed samples. In these simulations, the *Q ×* 1 population membership vector ***α****_j_* for individual *j* was sampled from a *Dirichlet*(***a***) distribution, where ***a*** is a vector containing ancestry weightings. In the most simple case ***a*** = *a***1**, where *a* is the admixture coefficient. For *a* = 0, this does not form a proper distribution and instead ancestry was selected by alternating individual ancestry between each of the populations. Increasing this coefficient increases admixture. When *a = 1,* this is effectively a uniform distribution and when *a* > 1, the mode of the distribution is one containing even admixture between all the populations.

The individual ancestries ***α****_j_* make up the rows of ancestry matrix ***A***, which has dimension *N × Q.* Multiplying this ancestry matrix by the population allele frequency vector (***P****_i_*), which (for a given SNP *i*) has length *Q,* generated an *N ×* 1 vector of allele frequencies for each individual 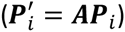 Individual genotypes *g_ij_* were generated from a 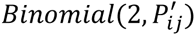 distribution.

To assess running time, the simulated datasets had *F_ST_* = 0.01, *M* = 100*k* SNPs and *N* ≈ {1*k*, 1.5*k*, 2*k*, 3*k*, 5*k*, 7*k*, 10*k*, 15*k*, 20*k*, 30*k*, 50*k*, 70*k*, 100*k*} individuals (since we used 6 populations of equal sample size, we rounded *N* to multiples of 6). Throughout this paper we report CPU time, but due to multi-threading present in the GSL^54^ and OpenBLAS libraries, run time was about 60% of CPU time. FastPCA accuracy was assessed using *M = 50k* SNPs and *N* ≈ *10k* individuals at *F_ST_* = {0.001,0.002,...,0.010}. Calibration and power of the selection statistic was assessed using 2 populations at *F_ST_* = {0.1,0.05,0.02,0.01,0.005,0.002,0.001,0.0005}, and also using 5 populations withthe tree structure described above. We set *M = 60k,* the effective number of independent SNPs in genotype array data^55^. When testing the power of the statistic, we wished to control the absolute difference in allele frequencies (*D*) between pairs of populations. For this purpose, SNPs under selection were generated in a similar manner as the above, except population allele frequencies were fixed at 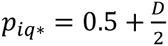 for one population and 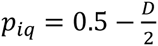 for the remaining population(s); this approximates allele frequency differences under a population genetic selection model with strong selection in one population, because the magnitude of allele frequency differences caused by strong selection is much larger than the magnitude of allele frequency differences caused by genetic drift.

### Assessing PC accuracy

Accuracy was assessed via the Mean of Explained Variances (MEV) of eigenvectors. Two different sets of *K N*-dimensional principal components each produce a *K*-dimensional column space. A metric for the performance of a PCA algorithm against some baseline is to see how much the column spaces overlap.

This is done by projecting the eigenvectors of one subspace onto the other and finding the mean lengths of the projected eigenvectors. If we have a reference set of PCs (***v***_1_, ***v***_2_,…,***v****_K_*) against which we wish evaluate the performance a set of computed PCs (*u*_1_*, u*_2_,..., *u*_K_), then the performance calculation becomes:

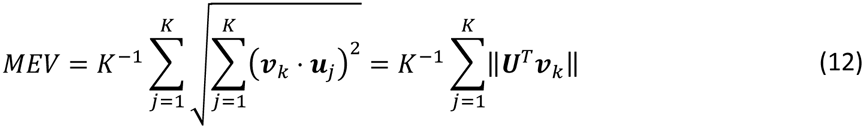

Here, ***U*** is a matrix whose column vectors are the PCs which we are testing. The test matrix can either be the result of another computation or the truth for a simulated sample. *K* eigenvectors can describe the population structure in a dataset with *K* + 1 populations. They can be constructed by first creating a vector 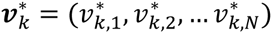 where 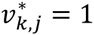 if individual *j* is in population *k* and 0 otherwise. The set of eigenvectors {***v***_1_, ***v***_2_,..., ***v****_K_*} are constructed by taking *K* of these vectors, normalizing them to have mean 0, and scaling/orthogonalizing them via the Gram-Schmidt process.

### GERA data set

The GERA dataset includes 62,318 individuals from Northern California typed on a European-specific 670,176-SNP array^56^. This dataset underwent two levels of filtration: a quality control step to produce the QC set of SNPs used to detect natural selection, and a second step used to produce the LD-pruned set of SNPs for PCA.

For the QC step, individuals were filtered to remove those with missing sex information, individuals related according to the provided pedigree data or with observed genomic relatedness greater than 0.05 in the GRM^58^ and individuals with less than 90% European ancestry as predicted by SNPweights^57^ using a worldwide dataset containing European, African, and Asian ancestry. After filtering, 54,734 individuals remained. Additionally, SNPs were initially filtered to remove non-autosomal SNPs, SNPs with minor allele frequency less than 1%, and SNPs with >1% missing data, leaving 608,981 SNPs.

The second stage of filtering removed SNPs that failed PLINK’s Hardy-Weinberg Equilibrium test^58^ with *p* < 10^−6^, and performed LD-pruning using PLINK. Due to regions of long-range LD, LD persisted even after one filtering run. Multiple rounds of LD filtering were performed using an *r*^2^ cutoff of 0.2 until additional rounds of LD filtering did not remove additional SNPs, leaving 162,335 SNPs.

FastPCA was run on the pruned set of 162,335 SNPs, while selection statistics were computed on the full set of 608,981 SNPs, prior to H-W filtering and LD-pruning. We note that many of the SNPs producing signals of selection generated significant H-W *p*-values (see Results - e.g. H-W *p* = 1.37 × 10^−79^ for LCT SNP rs6754311), which is an expected consequence of unusual population differentiation.

SNPweights^57^ was used to predict fractional Northwest European, Southeast European, and Ashkenazi Jewish ancestry for each individual. For plotting purposes, percentage ancestry in each of these three populations was mapped to an integer in [0,255], which was then used for the RGB color value for that sample, so a NW sample would appear red, SE would appear green and AJ would appear blue.

### PC Projection

POPRES^59^ individuals were projected onto these PCs. The left singular vectors (***U***) were generated by multiplying normalized genotypes for all SNPs in GERA (***Y****_GERA_*) by the PCs (***V***) and scaling by the singular values (**Σ**), the number of SNPs used to calculate the PCs (*M*) and the number of SNPs used for projection 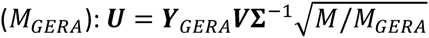. Projected PCs were then calculated by multiplying the corresponding set of SNPs in POPRES by these singular vectors and scaling again by the singular values: 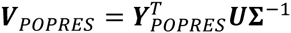 The projected individuals were overlaid on the PCA plot of GERA individuals and colored according to population membership and consistently with population assignment from SNPweights^57^.

## Results

### FastPCA Simulations

We used simulated data to compare the running time and memory usage of FastPCA to three previous algorithms: smartpca^26,31^, PLINK2-pca^58^, and flashpca^60^ (see Web Resources). We simulated genotype data from six populations with a star-shaped phylogeny using 100k SNPs (typical for real data after LD-pruning) and up to 100k individuals (see Methods). For each run, running time was capped at 100 hours and memory usage was capped at 40GB. The running time and memory usage of FastPCA scaled linearly with simulated dataset size (Figure 1), compared with quadratically or cubically for other methods. The computation became intractable at 50k-70k individuals for smartpca, PLINK2-pca and flashpca. The largest dataset, with 100k SNPs and 100k individuals, required only 56 minutes and 3.2GB of memory with FastPCA (Table S1). (We also note that shellfish (see Web Resources), a parallel PCA implementation, requires *O* (*MN*^2^ + *N*^3^) and is not computationally tractable on large data sets, as previously demonstrated^60^). Thus, FastPCA—unlike other publicly available software packages for analyzing genetic data—enables rapid principal components analysis without specialized computing facilities.

**Figure 1.**
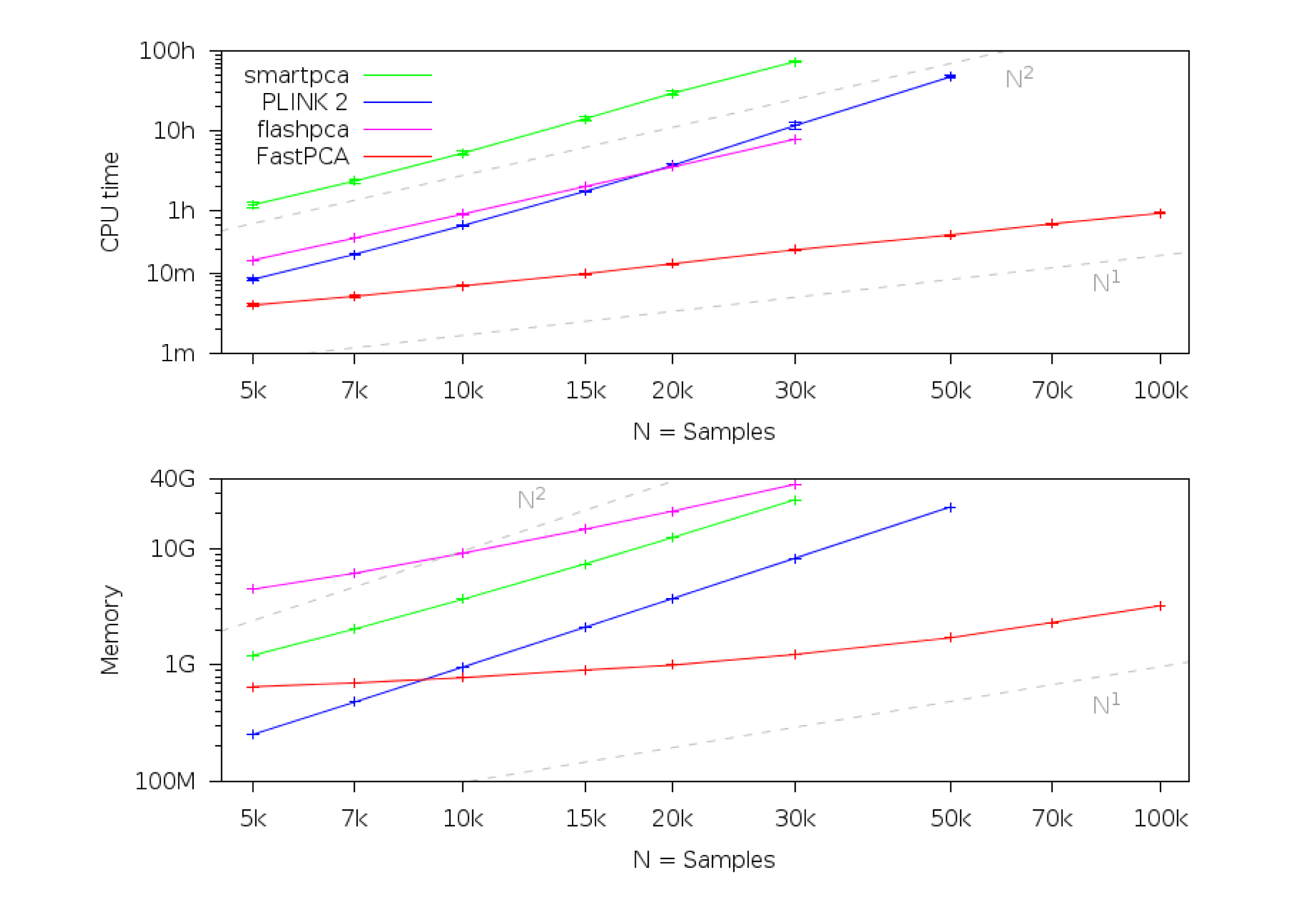
Running time and memory requirements of FastPCA and other algorithms. The CPU time and memory usage of FastPCA scale linearly with the number of individuals. On the other hand, smartpca and PLINK2-pca scale between quadratically and cubically, depending on whether computing the GRM (quadratic) or the eigendecomposition (cubic) is the rate-limiting step. The running time of flashpca scales quadratically (because it computes the GRM), but its memory usage scales linearly because it stores the normalized genotype matrix in memory. With 50k individuals, smartpca exceeded the time constraint (100 hours) and flashpca exceeded the memory constraint (40GB). With 70k individuals, PLINK2-pca exceeded the memory constraint (40GB). Run times are based on one core of a 2.26-GHz Intel Xeon L5640 processor; we caution that run time comparisons may vary by a small constant factor as a function of the computing environment. Numerical data are provided in Table S1.

We next assessed the accuracy of FastPCA, using PLINK2-pca^58^ as a benchmark. We used the same simulation framework as before, with 10k individuals (1,667k individuals per population) and 50k SNPs. We varied the divergence between populations, as quantified by *F_ST_* ^61^. We assessed accuracy using the Mean of Explained Variances (MEV) of the 5 population structure PCs (see Methods). We determined that the results of FastPCA and PLINK-pca were virtually identical (Figure 2). This indicates that FastPCA performs comparably to standard PCA algorithms while running much faster.

**Figure 2.**
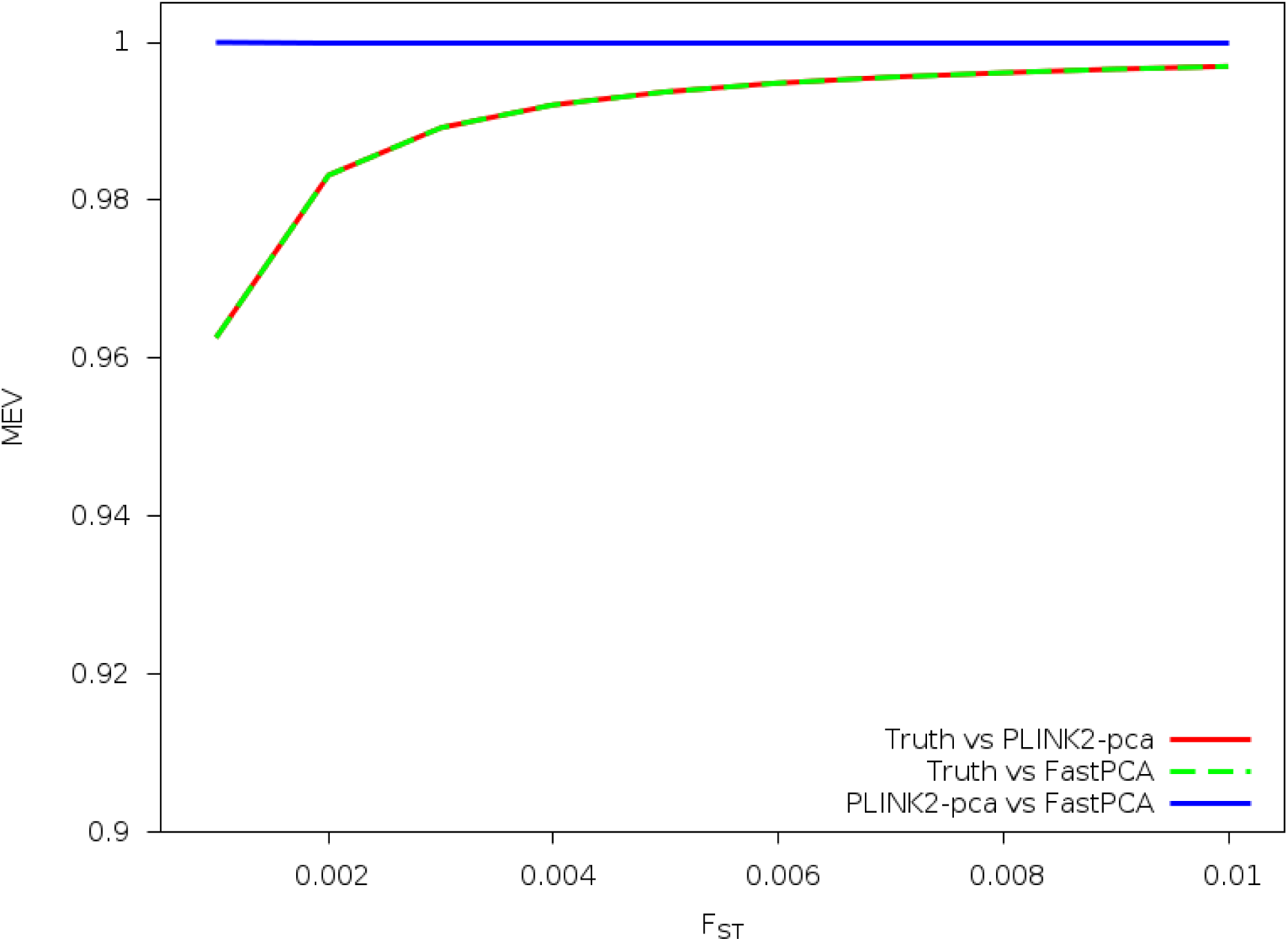
Accuracy of FastPCA and PLINK2-pca. FastPCA and PLINK2-pca were run on simulated populations of varying divergence. The simulated data comprised 50k SNPs and 10k total individuals from six subpopulations derived from a single ancestral population. PCs computed by PLINK2-pca and FastPCA were compared to the true population PCs and to each other using the Mean of Explained Variances (MEV) metric (see text). FastPCA explained the same amount of true population variance as PLINK2-pca in all experiments, and the methods output nearly identical PCs (MEV>0.999).

### PC-based Selection Statistic Simulations

We evaluated the calibration and power of the PC-based selection statistic. To evaluate calibration, we simulated 60k SNPs undergoing random drift with up to *N* = 50k individuals from two populations differentiated by *F_ST_* = {0.1,0.01,0.001}. At all values of *N* and *F_ST_*, the proportion of truly null SNPs reported as significant was well-calibrated at *p*-value thresholds ranging from 10*^−^*^1^ to10*^−^*^5^. Similar results indicating appropriate calibration were obtained for simulations with admixture (Table S2), as expected since the drift model still applies in the case of admixture^27^. The median of the selection statistic was slightly inflated at *F_ST_* = 0.1 due to a deficiency in the tail (Figure S1 and Table S2), but well-calibrated at the small values of *F_ST_* that correspond to our analyses of real data. The selection statistic in the presence of a population bottleneck performed identically to populations differentiated by the same *F_ST_* level (Table S2). We also simulated five populations with a phylogenetic structure (see Methods) that mimics the population structure found in the GERA data (see below) and found that the statistic remained well-calibrated here as well (Figure S1 and Table S2).

We evaluated power using the same number of SNPs and samples but at *F_ST_* = {0.1,0.05,0.02,0.01,0.005,0.002,0.001,0.0005} and using a separate set of SNPs under selection where the allele frequency between the two populations was varied (|D| = *|p*_1_ *− p_2_|*). The significance threshold was set to 8.3 × 10^−7^ based on 60K SNPs tested. There was no power to detect selection at *F_ST_* = 0.1. We observed a phase-change in the power simulations that was sharper for smaller *F_ST_*, where there was no power to detect selection below a specified allele frequency difference threshold, but there was complete power to detect selection at a slightly higher threshold (Figure 3a). We examined this effect in more depth using a range of samples sizes, and determined that the transition from no-power to complete-power was more sample size dependent at *F_ST_* = 0.001 (Figure 3b) than at *F_ST_* = 0.01 (Figure 3c), indicating that sample size is more important when analyzing more closely related populations. The PC-based selection statistic performed very similarly to the discrete-population test of selection^19^ in the case of data from discrete subpopulations (Figure S2). We also assessed effect of admixture on power by sampling ancestry for individuals between the two populations using a *Beta*(*a, a*) distribution. We determined that increasing the admixture parameter *a* (which reduces the variation in ancestry across samples) had a similar effect to reducing sample size (Figure S3).

**Figure 3.**
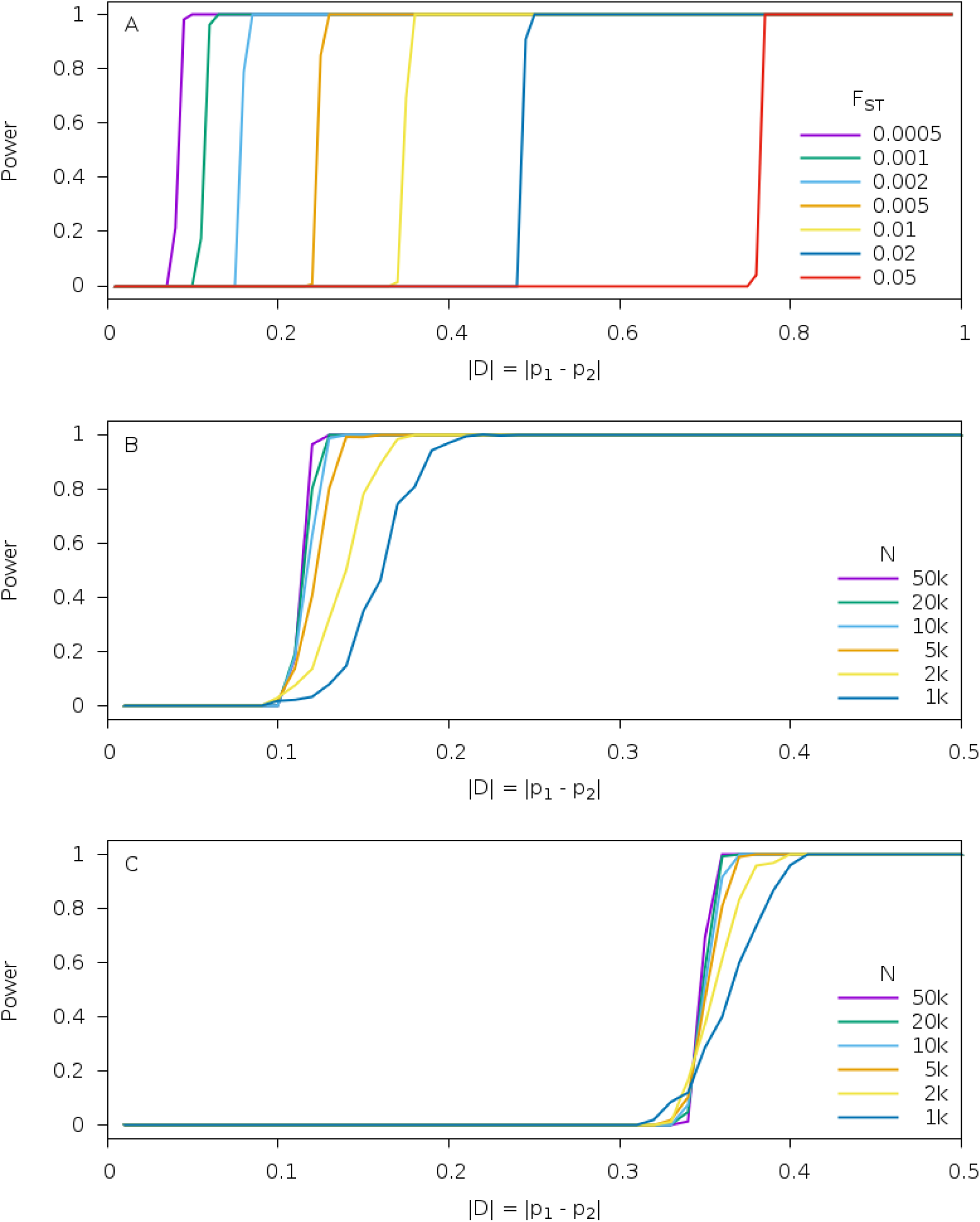
Power of PC-based selection statistic. The allele frequency difference at selected SNPs was varied between two populations separated by varying *F_ST_*. The significance threshold was set to 8.3 × 10^−7^ based on 60K SNPs tested. (a) With 50k samples, the power curves for *F_ST_* = {0.05,0.02,0.01,0.005,0.002,0.001,0.0005} showed a phase change. (b) Varying the number of samples for *F_ST_* = 0.001 demonstrated that this phase change was more gradual at smaller sample sizes. (c) Varying the number of samples at *F_ST_* = 0.01 showed that the impact of sample size was less pronounced than at *F_ST_* = 0.001.

### Application of FastPCA to a European American Cohort

We ran FastPCA on the GERA cohort (see Web Resources), a large European American dataset containing 54,734 individuals and 162,335 SNPs after QC filtering and LD-pruning (see Methods). This computation took 57 minutes and 2.6GB of RAM. PC1 and PC2 separated individuals along the canonical Northwest European (NW), Southeast European (SE) and Ashkenazi Jewish (AJ) axes^15^, as indicated by labeling the individuals by predicted fractional ancestry from SNPweights^57^ (Figure 4). These results are consistent with Banda *et al.* 2015^56^ which also examined this dataset. PC3 and PC4 detected additional population structure within the NW population.

**Figure 4.**
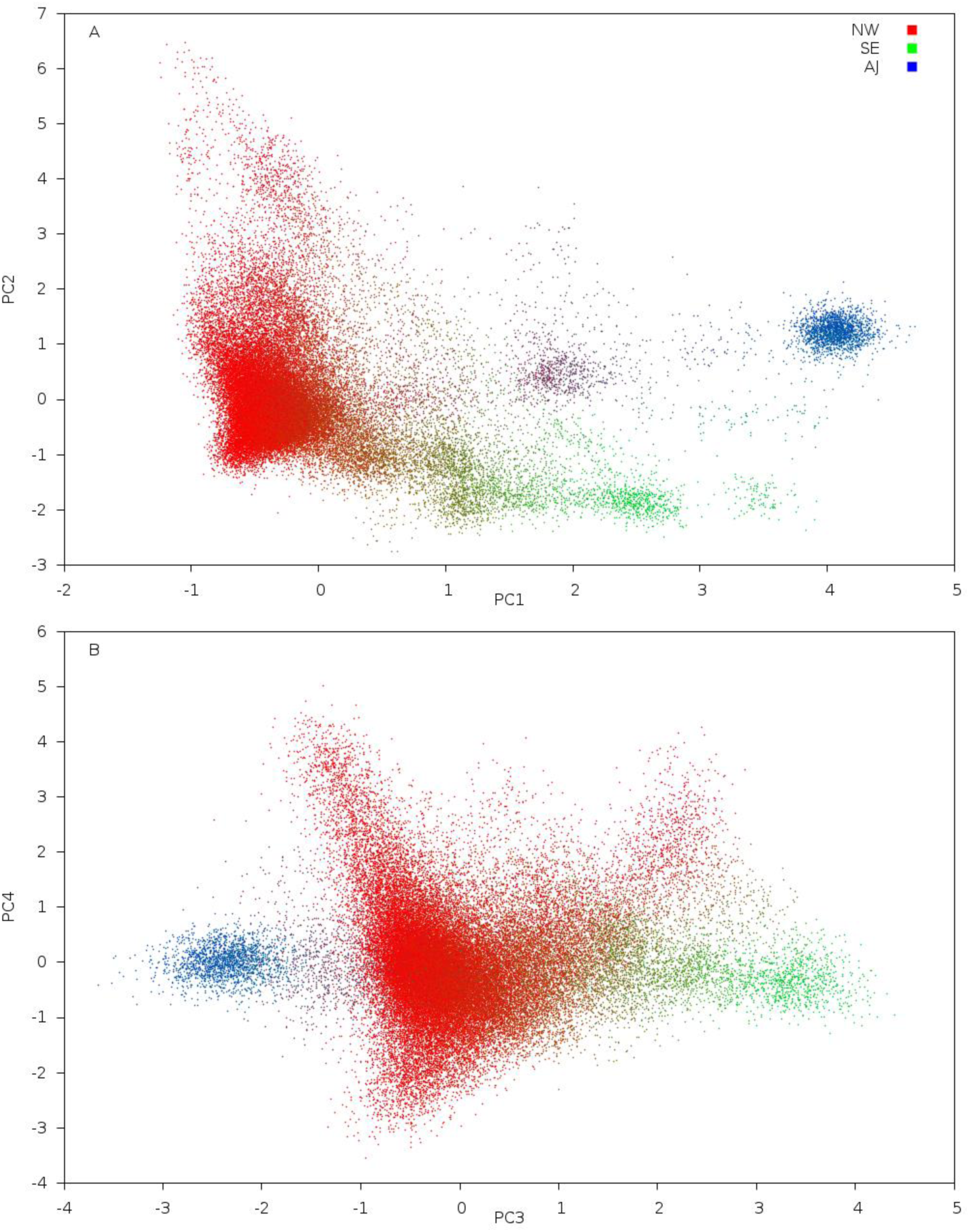
FastPCA results on GERA data set. FastPCA and SNPweights^57^ were run on the GERA cohort and the principal components from FastPCA were plotted. Individuals were colored by mapping Northwest European (NW), Southeast European (SE) and Ashkenazi Jewish (AJ) ancestry estimated by SNPweights to the red/green/blue color axes (see Methods). PC1 and PC2 separate the GERA cohort into northwest (NW), southeast (SE) and Ashkenazi Jewish (AJ) subpopulations. PC3 separates the AJ and SE individuals, while PC3 and PC4 further separates the NW European individuals.

To further investigate this subtle structure, we projected POPRES individuals from throughout Europe^59^ onto these PCs^31^ (see Methods). This analysis recapitulated the position of SE populations via the placement of the Italian individuals, and determined that PC3 and PC4 separate the NW individuals into Irish (IR), Eastern European (EE) and Northern European (NE) populations (Figure 5). This visual subpopulation clustering was confirmed via k-means clustering on the top 4 PCs, which consistently grouped the AJ, SE, NE, IR and EE populations separately (Figure S4. k-Means clustering confirms visually-observed subpopulations.Figure S4). We note that, in general, *K* PCs can cluster samples into *K*+1 subpopulations.

**Figure 5.**
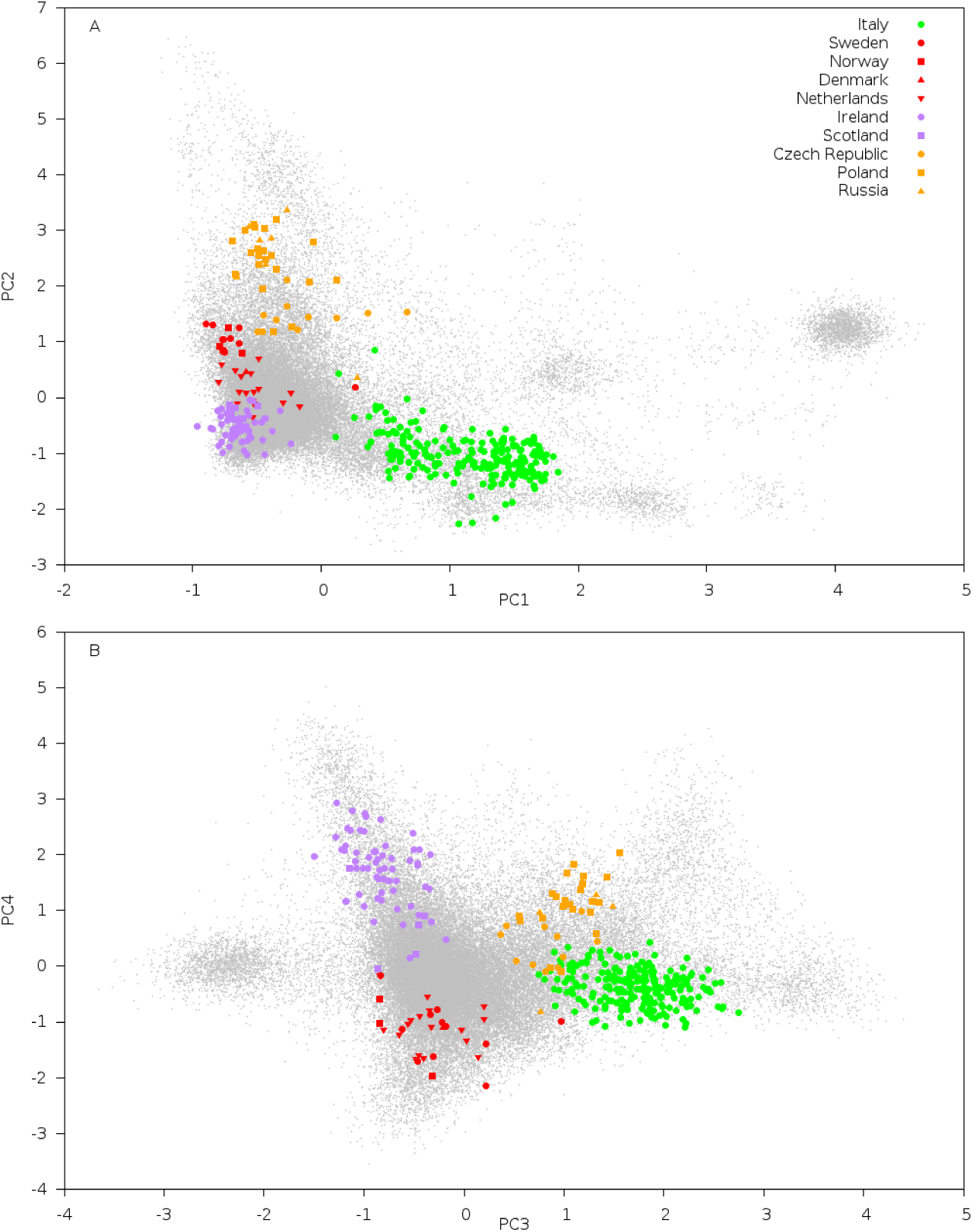
Separation of Irish, Eastern European and Northern European individuals in GERA data. We report results of projecting POPRES^59^ individuals onto top PCs. The plot of PC3 vs PC4 shows that the Northwest European (NW) individuals are further separated into Irish and Eastern European and Northern European populations. Projected populations were colored based on correspondence to the ancestry assignment from SNPweights^57^, except that Irish and Eastern European individuals were colored purple and orange, respectively, to indicate additional population structure.

### Application of PC-based Selection Statistic to a European American Cohort

For each of the top PCs, we computed our PC-based selection statistic for 608,981 non-LD-pruned SNPs (see Methods). The resulting Manhattan plots for PCs 1–4 are displayed in Figure 6 (QQ plots are displayed in Figure S5). Analyses of PCs 5–10 indicated that these PCs do not represent true population structure (Figure S6), but are either dominated by a small number of long-range LD loci^34,62,63^ or correlated with the missing data rate across individuals. Selection statistics for PCs 1–4 exhibited little or no inflation, particularly after removing Table 1 regions (Table S3).

**Table 1.**
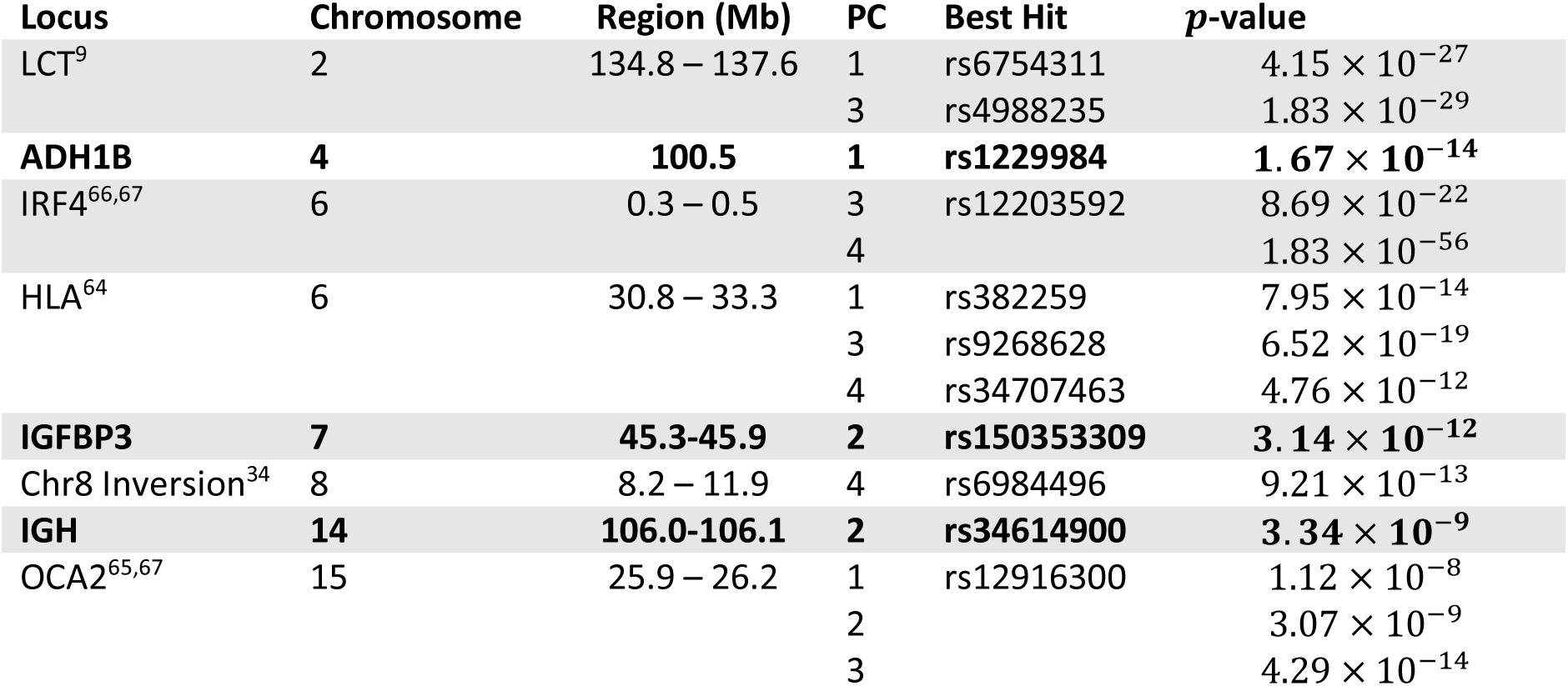
Genome-wide significant signals of selection in GERA data. We list regions with genome-wide significant (*α* = 0.05, Bonferroni correction with 608,981 SNPs × 4 PCs = 2,435,924 hypotheses tested, *p < 2.05 ×* 10*^−^*^8^) evidence of selection in the top 4 PCs. Loci that were not previously known to be under selection in Europeans are indicated in bold font. The chromosome 8 inversion signal is due to a PC artifact (see main text). Regions with suggestive evidence of selection (10*^−^*^6^ *< p < 2.05 ×* 10*^−^*^8^) are listed in Table S3.

**Figure 6.**
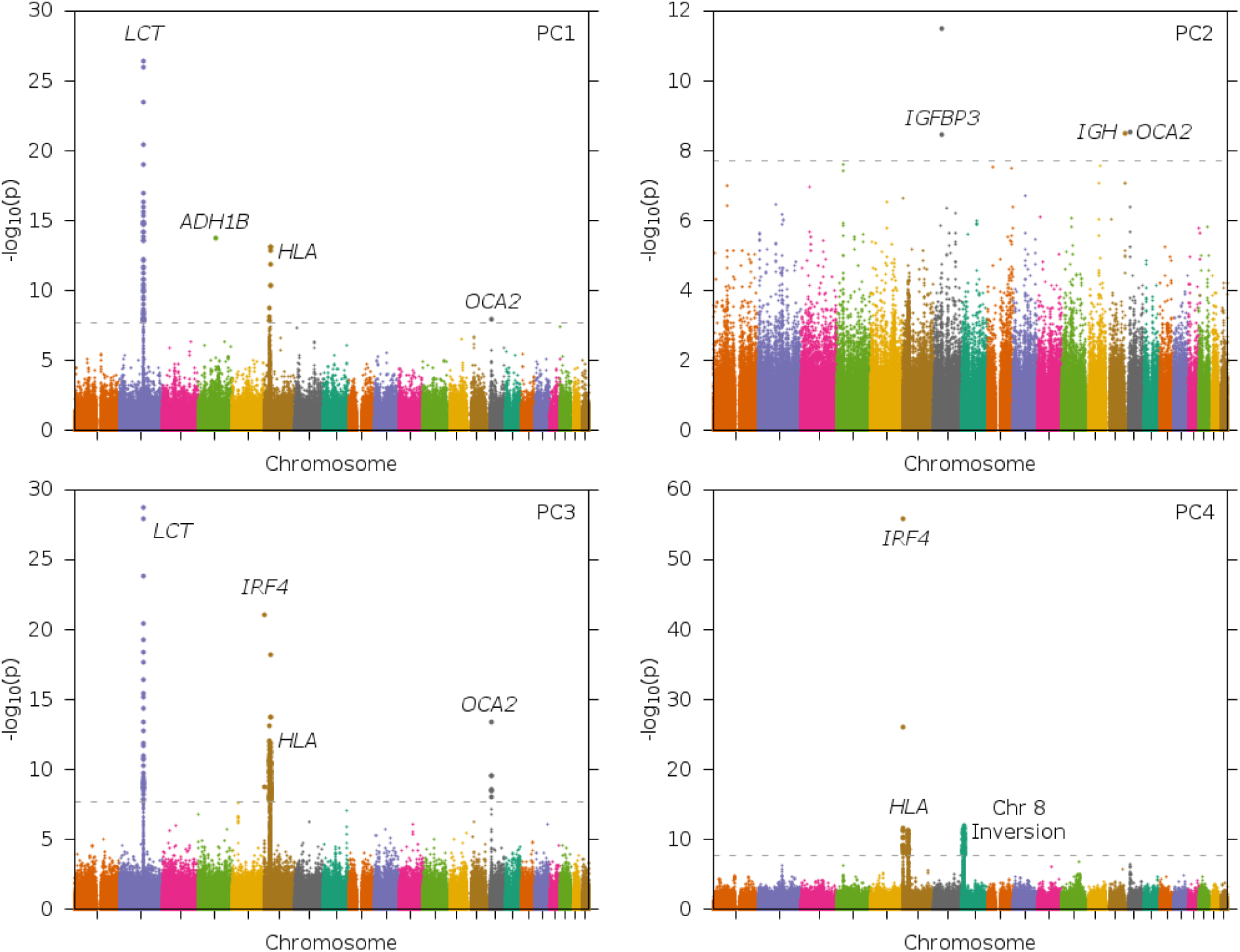
Signals of selection in the top PCs of GERA data. We display Manhattan plots for selection statistics computed using each of the top 4 PCs. The grey line indicates the genome-wide significance threshold of 2.05 × 10*^−^*^8^ based on 2,435,924 hypotheses tested (*α* = 0.05, 608,981 SNPs × 4 PCs).

Genome-wide significant signals (listed in Table 1) included several known selection regions^9,64–67^ and novel signals at ADH1B, IGFBP3 and IGH (see below). Suggestive signals were observed at additional known selection regions^66,68^ (Table S4). After removing the regions in Table 1, rerunning FastPCA and recalculating selection statistics, all of these regions remained significant except for a region on chromosome 8 with a known chromosomal inversion^34,62^ (Figure S7, Table S5). Thus, the remaining regions are not due to PC artifacts caused by SNPs inside these regions. We also found that a significantly greater proportion of SNPs under selection failed Hardy-Weinberg equilibrium, although the converse is not true, indicating that signals of selection are not a result of H-W artifacts (Figure S8). Detecting subtle signals of selection benefited from the large sample size, as subsampling the GERA data set at smaller sample sizes and recomputing PCs and selection statistics generally led to less significant signals (Table 2). We note that several suggestive selection signals, including signals at the known selected loci TLR1^66^ [MIM 601194] and SLC45A2^68^ [MIM 606202], are on the cusp of being significant and further increases to sample size may increase power to detect selection at suggestive loci.

**Table 2.** Performance of natural selection statistic in subsampled data. The selection statistic was computed in random subsets of individuals of specified size for each SNP in Table 1 (except for the chromosome 8 inversion region) and the known selection regions TLR1^66^ and SLC45A2^68^ in Table S4. We report the median selection statistic P-value across 100 random subsets.

| Locus | SNP | Sample size |  |  |  |  |  |  |
| --- | --- | --- | --- | --- | --- | --- | --- | --- |
|  |  | Full data set | 1k | 2k | 5k | 10k | 20k | 50k |
| LCT | rs6754311 | 2.15e-25 | 4.91e-17 | 2.97e-20 | 1.53e-23 | 1.17e-24 | 2.63e-25 | 1.02e-26 |
|  | rs4988235 | 1.15e-27 | 7.44e-17 | 9.80e-20 | 4.64e-23 | 3.11e-24 | 2.69e-25 | 1.62e-27 |
|  | rs17346504 | 8.41e-7 | 2.86e-2 | 1.25e-2 | 9.49e-4 | 6.03e-5 | 8.12e-6 | 9.80e-7 |
| ADH1B | rs1229984 | 1.26e-13 | 3.91e-9 | 3.51e-11 | 1.97e-12 | 5.54e-13 | 1.50e-13 | 1.31e-13 |
| IRF4 | rs12203592 | 5.52e-55 | 3.15e-6 | 9.18e-12 | 7.47e-25 | 7.21e-36 | 7.02e-45 | 2.19e-54 |
| HLA | rs382259 | 5.38e-13 | 8.68e-9 | 1.23e-10 | 7.07e-12 | 1.85e-12 | 7.51e-13 | 5.77e-13 |
|  | rs9268628 | 8.66e-18 | 3.62e-5 | 3.41e-7 | 5.97e-12 | 2.10e-14 | 2.68e-16 | 1.00e-17 |
|  | rs4394275 | 9.36e-12 | 8.40e-2 | 1.94e-3 | 1.44e-5 | 4.00e-8 | 7.86e-10 | 1.24e-11 |
| IGFBP3 | rs150353309 | 5.82e-12 | 5.90e-4 | 1.49e-5 | 2.72e-8 | 3.61e-10 | 3.34e-11 | 6.61e-12 |
| IGH | rs34614900 | 5.23e-9 | 6.33e-3 | 2.24e-4 | 2.26e-6 | 2.01e-7 | 3.32e-8 | 5.32e-9 |
| OCA2 | rs12916300 | 2.80e-13 | 6.29e-6 | 1.07e-7 | 3.67e-9 | 1.94e-11 | 5.29e-12 | 3.11e-13 |
|  | rs2703951 | 5.11e-7 | 1.12e-1 | 2.45e-2 | 7.96e-4 | 7.17e-5 | 4.52e-6 | 5.74e-7 |
| TLR1 | rs5743611 | 5.42e-8 | 8.05e-3 | 4.27e-4 | 9.41e-6 | 1.19e-6 | 2.17e-7 | 5.60e-8 |
|  | rs4833095 | 6.52e-7 | 6.07e-4 | 3.37e-4 | 7.35e-5 | 3.64e-5 | 6.03e-6 | 7.10e-7 |
| SLC45A2 | rs16891982 | 6.89e-8 | 8.25e-4 | 2.17e-4 | 1.93e-5 | 4.55e-6 | 2.46e-7 | 7.31e-8 |

We identified a genome-wide significant signal of selection at rs1229984, a coding SNP (Arg47His) in the ADH1B alcohol dehydrogenase gene (Table 1). The allele rs1229984*T has been shown to have a protective effect on alcoholism risk^44–47^ and to produce an REHH signal in East Asians^14,46,48,49^, but was not previously known to be under selection in Europeans. (Previous studies noted the higher frequency of the rs1229984*T allele in western Asia compared to Europe, but indicated that selection or random drift were both plausible explanations^69,70^.) We examined the allele frequency of the rs1229984*T allele in the five subpopulations: AJ, SE, NE, IR and EE (Table S7). We observed allele frequencies of 0.21 in AJ, 0.10 in SE, and 0.05 or lower in other subpopulations, consistent with the higher frequency of the rs1229984*T in western Asia. A comparison of NE to the remaining subpopulations using the discrete subpopulation selection statistic^19^ also produced a genome-wide significant signal after correcting for all hypotheses tested (Table S7); this is not an independent experiment, but indicates that this finding is not due to assay artifacts affecting PCs.

To further understand the selection at this locus, we examined the allele frequency of rs1229984*T in 1000 Genomes project^71^ populations (see Web Resources), along with the allele frequency of the regulatory SNP rs3811801 that may also have been a target of selection in Asian populations^46^. The haplotype carrying rs3811801*A (and corresponding haplotype H7) was absent in populations outside of East Asia (Table S8). This indicates that if natural selection acted on this SNP in Asian populations, selection acted independently at this locus in Europeans. One possible explanation for these findings is that rs1229984 is an older SNP under selection in Europeans, while rs3811801 is a newer SNP under strong selection in Asian populations leading to the common haplotype found in those populations.

The IGFBP3 insulin-like growth factor-binding protein gene had two SNPs reaching genome-wide significance. Genetic variation in IGFBP3 has been associated with breast cancer^72^, height^73^, blood pressure^74^ and hypertension^75^, although the published associated SNPs are not in LD with the two SNPs we detected. The IGH immunoglobulin heavy locus had one genome-wide-significant SNP and two suggestive SNPs with *p*-value < 10^−6^ (Table 1). Genetic variation in IGH has been associated with multiple sclerosis^76^, although the published associated SNPs are not in LD with the three SNPs we detected. The IGFBP3 and IGH SNPs each had substantially higher minor allele frequencies in Eastern Europeans, but were not genome-wide significant under the discrete subpopulation selection statistic^19^(Table S9 andTable S10). The existence of multiple SNPs at each of these loci with p < 10^−6^ for the PC-based selection statistic suggests that these findings are not the result of assay artifacts.

## Discussion

We have detected new, genome-wide significant signals of selection by applying a PC-based selection statistic to top PCs computed using FastPCA, a computationally efficient (linear-time and linear-memory) algorithm. Although mixed model association methods are increasingly appealing for conducting genetic association studies^55,77^, we anticipate that PCA will continue to prove useful in population genetic studies, in characterizing population stratification when present in association studies, in supplementing mixed model association methods by including PCs as fixed effects in studies with extreme stratification, and in correcting for stratification in analyses of components of heritability^78,79^. Our PC-based selection statistic extends previous statistics developed for discrete populations^19^. In contrast to previous work on detecting selection using PCs^63,80^ or using the spatial ancestry analysis (SPA) method^81^, our statistic is able to detect signals at genome-wide significance, a key consideration in genome scans for selection^82^. Our work demonstrates the advantages of comparing closely related populations in very large sample sizes to detect novel loci, whereas very recent studies applying related methods to smaller sample sizes detected genome-wide significant signals only at previously known loci^83,84^. In particular, we detected genome-wide significant evidence of selection in Europeans at the ADH1B locus, which was previously reported to be under selection in East Asian populations^14,46,48,49^ using REHH^53^ (which can only detect relatively recent signals and does not work on standing variation^3^). We also detected genome-wide significant evidence of selection at the disease-associated IGFBP3 and IGH loci. While the SNPs under selection at these loci are not in LD with the disease-associated SNPs identified in previous association studies, these genes are biologically important and there may be other phenotypes associated with the selected SNPs. Although we emphasize the importance of genome-wide significance, loci with suggestive signals of selection that do not reach genome-wide significance could potentially be used to increase the power of disease mapping^85^.

We note that our work has several limitations. First, top PCs do not always reflect population structure, but may instead reflect assay artifacts^86^ or regions of long-range LD^34^; however, PCs 1–4 in GERA data reflect true population structure and not assay artifacts, because the PCs (and the signals of selection they detect) remained nearly unchanged after removing regions with significant signals of selection (Table 1) and rerunning PCA. Second, common variation may not provide a complete description of population structure, which may be different for rare variants^87^; we note that based on analysis of real sequencing data with known structure, we recommend that LD-pruning and removal of singletons (but not all rare variants) be applied in data sets with pervasive LD and large numbers of rare variants (see Appendix). Third, our selection statistic is only capable of detecting that selection occurred, but not when or where it occurred; indeed, top PCs may not perfectly represent the geographic regions in which selection occurred, underscoring that interpretation of results can be a fundamental limitation of model-free methods. Fourth, our selection statistic performs best when allele frequencies vary linearly along a PC; the SPA method^81^ (see above) models allele frequency as a logistic function and is not constrained by this limitation. Despite these limitations, we anticipate that FastPCA and our PC-based selection statistic will prove valuable in analyzing the very large data sets of the future.

## Appendix

Inferring ancestry from genetic data is a common problem in both population and medical genetic studies, and many methods exist to address it^31,32,88^. Principal components analysis (PCA)^31^ has been shown to be effective at elucidating geographic structure from genetic data^89^ and correcting for confounding due to population stratification in association mapping^26^. These uses of PCA depend critically on its ability to separate genetically disparate subpopulations when analyzing data from commercial genotyping arrays. However, as high-throughput sequence data becomes more common, enabling ancestry inference from this new class of data is becoming increasingly relevant.

As sequence data contains more variants, and many more population-specific variants^90^, it may be reasonable to expect that PCA applied to high-throughput sequence data will be substantially more effective than the corresponding analysis on genotype data. However, our results suggest the opposite. Specifically, PCA makes assumptions about marker independence that are violated by the pervasive linkage disequilibrium in sequence data. In addition, assumptions about genetic drift that are reasonable for common SNPs on genotyping arrays are less so when applied to the numerous rare variants in sequence data^87^.

### Methods

Principal Components Analysis (PCA) is generally applied to a genetic relationship matrix (GRM) that is computed as:

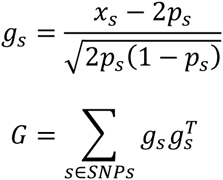

where *x_s_* is a vector of genotypes for SNP s and *p_s_* is the minor allele frequency of SNP *s.* We propose modifications to standard PCA to deal with two challenges that are present in sequence data but absent from genotype data: pervasive linkage disequilibrium, and rare variants. Specifically, we recommend that LD pruning be applied to sequence data and singleton variants be removed. While we evaluated more sophisticated approaches to handling these issues, they did not improve our results beyond these simpler approaches. Importantly, we recommend against a commonly used strategy of removing all low frequency of rare variants as these variants contain significant information for detecting population structure.

### Linkage Disequilibrium

It is well known that application of PCA to regions of the genome containing long-range LD blocks can confound PCA’s ability to separate disparate populations^31,63^. As a result, these LD blocks are often simply excluded from analysis. However, in sequence data, many regions of the genome outside of previously identified long-range LD blocks contain sufficient LD to bias results. As a result, we examine three methods to deal with LD: (1) LD Pruning (2) LD Shrinkage^63^ and (3) LD Regression^31,9,12,9^.

LD Pruning is a commonly applied approach to removing correlated SNPs from a dataset. To produce a data set pruned for LD above a threshold T, one SNP of any pair of SNPs in LD (*r*^2^ *>* T) is removed from the data.

LD Shrinkage is a more sophisticated method of correcting for LD proposed by (Zou et al. 2012)^63^. In LD shrinkage, each SNP s is weighted by its LD to surrounding SNPs before inclusion in the genetic relationship matrix.

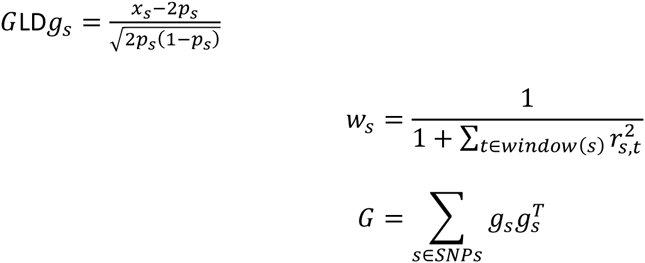

We note that *t ∈ window*(*s*) refers to SNPs *t* that are within some region of the genome surrounding SNP *s.* Intuitively, this is a heuristic to correct for the over representation in the GRM of some SNPs that are redundant with respect to nearby SNPs.

LD Regression was originally proposed in (Patterson et al. 2006)^31^ and utilized extensively in (Gusev et al. 2013) ^91^. Only the residual of a SNP—after regressing out other SNPs in LD—in the GRM.:

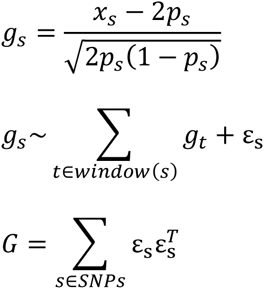

### Rare Variants

In considering how to optimally include rare variants in the genome, we examined three strategies. The first strategy was to include all rare variants as described in the computations above without any modifications. The second strategy was to exclude all variants below a threshold, which is a standard strategy used in several recent papers. We compared these simple strategies to a strategy based on reweighting rare variants to optimize the separation between populations.

We considered a particular scenario to optimize. Specifically, we imagine that two populations that split from one another *t* generations ago are equally represented in our GRM. We would like to optimize the proportion of variance in our GRM that is explained by the true population labels. That is, our figure of merit is:

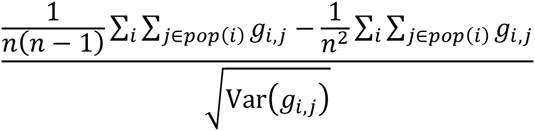

where *pop*(*i*) refers to the subpopulation from which individual *i* came.

Now, considering the population split, our data contains two classes of variants: those variants that are result of mutations predating the population split (pre-split SNPs), and those variants arising after the population split (post-split SNPs). For pre-split SNPs we invoke the normal approximation to genetic drift described. That is, the difference between allele frequencies *p*_1_*, p_2_* (for populations 1 and 2, respectively) is:

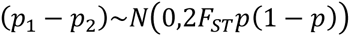

where *p* is the allele frequency in the ancestral population prior to the split and *F_ST_* quantifies the genetic drift that has occurred since the split. We note that this approximation is reasonable for common SNPs and for small values *F_ST_*. If we assume that our data contains only pre-split SNPs then our figure of merit is optimized by the standard computation of the GRM given above. On the other hand, rare, post-split SNPs have the property that

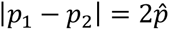

where 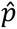 is the allele frequency estimated from the sample. This difference implies that the optimal weighting for pre-split SNPs is 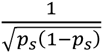 identically:

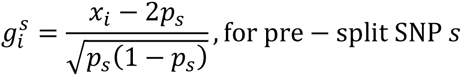

but the optimal weighting for post-spit SNPs is 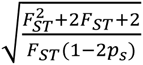

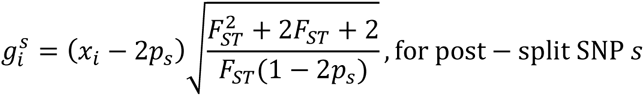

However, this modification requires knowledge of the *F_ST_* between studied subpopulations and, more dauntingly, which SNPs are post-split. We believe it is reasonable to iterate over several values of *F_ST_* (and find that in real data results are relatively robust to choice of *F_ST_*). In order to deal with uncertainty over the set of post-split SNPs, we propose that a SNP be considered post-split if

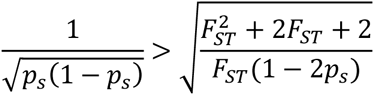

We examine the effect of both of these modifications on the effectiveness of PCA to separate genetically disparate subpopulations.

### Analysis of Northern vs. Southern Europe in POPRES Targeted Sequencing Data

We analyzed 531 individuals from the UK referred to as Northern European and 146 Italian, 134 Portuguese, 100 Spaniards, and 7 Swiss Italian individuals collectively referred to as Southern European^10^. We excluded 25.9 kb of sequence data from genes on the × chromosome, focusing solely on the autosomes. In total, 8,469 SNPs were polymorphic in either of the Northern or Southern European Samples. These variants were overwhelmingly rare, with 81.5% of variants having a MAF < 1% in the combined sample.

We tested various methods to correct for LD and better handle rare variants (see Methods). The results are summarized in Table S11. These results indicate that handling of both rare variants and LD is critical to maximizing the performance of PCA on this class of data. Applying standard PCA, the top 5 PCs explained only 2.3% of the variance (r^2^=0.023) of the true population labels. This was improved substantially by removing or reweighting rare variants with (r^2^=0.287, 0.341, 0.352) for removing variants with MAF < 0.02, removing singletons and reweighting, respectively. This indicates that rare variants, particularly singletons, may be problematic when analyzed using PCA. However, the difference between removing variants with MAF < 0.02 and reweighting (r^2^=0.287 vs 0.352) suggests that these variants do contain useful information for ancestry inference and should not be universally excluded.

Additionally, application of a method to correct for LD significantly improved performance of PCA when performed in conjunction with singleton exclusion or rare variant reweighting. With rare variant reweighting, LD shrinkage ^8^ (r^2^=0.563) performing slightly better than LD regression (r^2^=0.528) ^2^ and LD runing (r^2^=0.534). While LD Pruning performed well, this may be due to the fact that LD is broken up because the dataset contains sequence data from separated chunks of genome.

### Recommendations

In data sets that do not include pervasive LD or large numbers of rare variants (i.e. genotyping data), standard techniques are likely to be successful in detecting population structure. However, in data sets that have pervasive LD and large numbers of rare variants, we recommend that LD pruning and singleton removal be applied. While more sophisticated methods for dealing with these issues were assessed, we did not observe significant improvements above and beyond these simpler approaches. Importantly, we do not recommend that all low frequency and rare variants (MAF < 0.02) be removed as these variants do significantly improve detection of population structure.

## Acknowledgements

We are grateful to D. Reich for helpful discussions and S. Pollack for assistance with FastPCA software. This research was funded by NIH grant R01 HG006399. SM is funded by NSF grants DMS-1209155 and DMS-1418261.

## Web Resources

EIGENSOFT version 6.1, including open-source implementation of FastPCA and smartpca and the PC-based selection statistic: https://data.broadinstitute.org/alkesgroup/EIG6.1/

PLINK2: https://www.cog-genomics.org/plink2/

flashpca: https://github.com/gabraham/flashpca

GERA cohort: http://www.ncbi.nlm.nih.gov/projects/gap/cgi-bin/study.cgi?study_id=phs000674.v1.p1

1000 Genomes: http://www.1000genomes.org/

Shellfish: http://www.stats.ox.ac.uk/~davison/software/shellfish/shellfish.ph

### Description of Supplemental Data

Supplemental Data include eight figures and eleven tables.

